# Fine Social Discrimination of Siblings in Mice: Implications for Early Detection of Alzheimer’s Disease

**DOI:** 10.1101/2024.10.14.617786

**Authors:** Lola MP Fauré, Sébastien Gauzin, Camille Lejards, Claire Rampon, Laure Verret

**Affiliations:** Centre de Recherches sur la Cognition Animale, Centre de Biologie Intégrative, Université de Toulouse; CNRS, UPS, 31062, France

## Abstract

The ability to distinguish between individuals is crucial for social species and supports behaviors such as reproduction, hierarchy formation, and cooperation. In rodents, social discrimination relies on memory and the recognition of individual-specific cues, known as "individual signatures". While olfactory signals are central, other sensory cues —such as auditory, visual, and tactile inputs — also play a role. However, little research has explored the fine discrimination of individuals with overlapping cues, such as siblings or cohabitating mice. This study investigates whether mice can discriminate between two closely related individuals: siblings from the same litter and cage. We tested the hypothesis that it would be more challenging for mice to distinguish between siblings than between unrelated mice due to shared cues. Moreover, social cognitive impairments are common in neurodegenerative diseases like Alzheimer’s disease (AD), where difficulties in recognizing faces and voices progressively disrupt social interactions in patients. Using a mouse model of AD (Tg2576), known for the progressive onset of cognitive deficits, we assessed whether the ability to discriminate between siblings is preserved in “pre-symptomatic” animals. Thus, we first demonstrated that male and female C57BL6/J mice can discriminate siblings, regardless of sex. Next, we revealed that “pre-symptomatic” 3-month-old Tg2576 mice exhibit impairments in fine social memory, while their general social memory remains unaffected. Thus, we demonstrate that the inability to perform fine social discrimination is an early cognitive impairment that arises before other well-documented memory abnormalities in this AD mouse model.

**SIGNIFICANCE STATEMENT:** Discriminating between individuals is vital for social species, supporting reproduction, hierarchy formation and cooperation. This study examined whether mice can perform fine social discrimination, *i.e.*, distinguish between siblings from the same home cage. We found that both male and female C57BL6/J mice can discriminate siblings, regardless of their sex. Notably, this fine social discrimination ability is impaired in mouse model of Alzheimer’s disease (AD) at a very early stage of the pathology, *i.e.,* at an age when they can still recognize two unrelated mice from different home cages. This suggests that fine social discrimination deficits occur early in AD, and could serve as an early marker of the pathology.

## INTRODUCTION

In animals, the ability to distinguish between individuals allows the identification of familiar and unfamiliar individuals and is therefore crucial for social animal species. In rodents, for instance, social discrimination enables appropriate social interactions, such as reproduction, cooperative behavior, the establishment of dominance hierarchies, aggression, and avoidance (Okuyama, 2018), all of which are of primary importance for species survival. The capacity to distinguish between two individuals is largely based on social discrimination memory, which is the ability to remember and identify known conspecifics (Wang and Zhan, 2022). Individual discrimination is a more complex process, embedded within the broader concept of social discrimination. It involves recognizing specific characteristics such as group membership, kinship, age, sex, reproductive status, or hierarchical position by taking into account idiosyncratic traits (Gheusi et al., 1997). Additionally, each individual carries an “individual signature” that facilitates social discrimination. This signature is defined as a set of complex phenotypic traits that are individually distinctive to some extent, meaning they exhibit greater inter-individual variation than intra-individual variation (Beecher,1982). Furthermore, Hepper (1991) suggests that the ability to discriminate between individuals is based on the similarity of the “signatures” of relatives, which is closely linked to their genotype. Signatures of phenotypic variation derived from an individual’s genome are also influenced by environmental factors like bacterial flora and diet, which can affect body odor (Halpin, 1986). According to Brown (1979), rodents recognize their conspecifics and learn about their social surroundings primarily through olfactory signals, including pheromones, urine, and body odor. However, many studies suggest that the individual signature enabling individual discrimination is not solely based on olfaction but also includes auditory, tactile, and visual cues (Brudzynski, 2013; Simola and Granon, 2019; Arakawa, 2020; Jabarin et al., 2022). It has also shown that facial expressions in rodents may convey information about their emotional state. For instance, mice react differently to painful stimuli, notably by puffing up their cheeks and noses and changing the position of their ears and whiskers (Langford et al., 2010). This supports the idea that social discrimination relies on multifactorial cues.

However, the fine discrimination ability of mice in situations where individual cues overlap, such as genetic similarity in the case of siblings or shared social odors in cohabitating mice, has received limited attention.

Many brain illnesses, such as autism, schizophrenia, and Alzheimer’s disease (AD), are associated with social cognitive impairments (Kennedy and Adolphs, 2012; Henry et al., 2016). In AD, these deficits appear progressively and result in an inability to distinguish emotions from facial expressions (Hargrave et al., 2002; Bediou et al., 2009), voices (Roberts et al., 1996), or body movements (Koff et al., 1999). These impairments can interfere with social communication (Shimokawa et al., 2001), leading to difficulties in maintaining interpersonal relationships. Due to the multifactorial aspect of social discrimination, social cognitive alterations may be subtle and difficult to identify in the early stages of AD (Henry et al., 2016). However, early detection of social cognition deficits would enable better patient management, aiming to limit or slow the progression of symptoms.

In this study, we first investigated whether a healthy mouse can accurately discriminate between two individuals with whom she is not related, but who are closely related to each other, such as brothers or sisters. We hypothesized that two siblings of the same age and sex, born to the same parents, from the same litter, and living in the same home cage, would be more difficult to discriminate than two unrelated mice born to different parents and raised in separated cages. We then examined whether this type of discrimination was preserved at an early stage of Alzheimer’s disease in the Tg2576 mouse model, known for developing progressive cognitive deficits.

## METHODS

### Animals

All mice were housed in groups of 2 to 5 per cage, kept on a 12-hour light/12-hour dark cycle, with free access to food and water. All animals were handled an average of three times a week before behavioral experiments. Mice displaying signs of barbering were excluded from the study. All experiments were conducted in accordance with European Union guidelines (2010/63/EU) for the care and use of laboratory animals. Our animal facility is fully accredited by the French Direction of Veterinary Services (E 31-555-011, June 17^th^, 2021), and experimental procedures conducted in this study were authorized by local ethical committees and the French Ministry for Research (2023100310155057_v4#44099).

### C57BL6/J mice

Social memory assessments were performed using male (n=23) and female (n=21) C57BL/6J mice (WT; Charles River, France), aged 3 to 4 months. Males (n=12) and females (n=21) were tested in the five-trial test, followed by the three-chamber test. During these assays, the tested-mice encountered male or female demonstrators. An additional group of males (n=11) was used for the seven-trial test, during which the tested-mice encountered female demonstrators.

### Tg2576 mice

Social memory was also assessed in the Tg2576 transgenic mouse model of AD (Hsiao et al., 1996) by an experimenter blind to the genotype. Female Tg2576 mice and their non-transgenic littermates (NTg) were bred and genotyped in-house as described previously (Verret et al., 2013). Three-month-old NTg (n=7) and Tg2576 (n=9) mice underwent the five-trial test, which included a fine social discrimination test (trial 5: presentation of two sisters) followed by a classic social discrimination test (trial 5: presentation of two unrelated individuals from different cages). All demonstrators were NTg females.

### Demonstrators

Across all experiments, the demonstrators were brothers or sisters aged 4 to 12 months and housed together after weaning. Demonstrators and tested mice were unrelated (from different parents) and had never encountered each other before the tests. Mice with deteriorated fur or whiskers were excluded from the procedure.

### Behavioral experiments

In all behavioral tests (5-trial social memory, sociability and social discrimination tests, and 7-trial test), C57BL6/J females were exposed to female (n=6) or male (n=12) demonstrators, while C57BL6/J males were exposed to male (n=6) or female (n=6) demonstrators. Females from the Tg2576 line (NTg, n=7; Tg2576, n=10) were tested in the 5-trial test and interacted with NTg sisters. The duration of social interactions was scored in real-time using the Ethoc software developed in-house. A social interaction was defined as the tested mouse being within 1 cm of the demonstrator, oriented toward it, and engaging behaviors like sniffing or attempting to grab it with its paws. Between each trial, all equipment and testing arena were cleaned with 30% ethanol. The same mice first underwent the 5-trial social memory test, followed by the 3-chamber test.

### Five-trial social memory test

This test assesses short-term memory for social discrimination. The tested mouse was placed for 10 min in a 48 × 37 × 21 cm arena containing a small circular cage with bars (Domínguez et al., 2019). An unfamiliar, unrelated demonstrator mouse [a] was introduced into the cage for four successive 5-minute, with10-minute intervals between trials. On the fifth trial, a novel demonstrator mouse [a’] — a sister of the previous demonstrator, now considered familiar — was presented. Both demonstrators were from the same litter and housed in the same cage after weaning.

### Sociability and social discrimination tests

Mice were placed in the center of a three-chamber setup (Moy et al., 2004) and allowed to explore for 10 min. To assess sociability, the tested mouse was introduced to an unrelated mouse [a], confined in a small cylindrical cage. On the following day, the tested mouse was introduced to a new mouse [b], from a different litter and cage, with the previously encountered demonstrator mouse [a] in the other chamber, now considered familiar. On the final day, fine social discrimination was assessed by introducing two siblings (either sisters or brothers) raised in the same cage. One was the familiar mouse (used as demonstrator on days 1 and 2, mouse [a]), and the other was a previously unknown sibling of [a] (mouse [a’]).

### Seven-trial test

The 7-trial test, developed by Loisy et al. (2022), evaluates long-term social discrimination memory. On the first day, mice were placed in a 48 × 37 × 21 cm arena containing a small circular cage with metal bars for 10 min. An unfamiliar, unrelated mouse [a] was introduced for four successive 5-minute trials, with 10-minute intervals between trials. On the fifth trial, a novel demonstrator mouse [b] was presented. On the second day, mice were split into two groups and subjected to one of the following conditions:

- Option 1: the same demonstrator mouse [a] from day 1 was presented twice, assessing social memory of the original mouse.
- Option 2: a sister [a’] of the day 1 demonstrator [a] was presented twice. This allows to evaluate fine social discrimination over the long term.

For both options, demonstrators were presented for two 5-minute trials, following the same procedure as day 1.

### Statistical analysis

Statistical analyses were performed using Prism 9. Repeated measures (RM) two-way ANOVA were used to evaluate trial effects and demonstrator genotype or sex effects in the 5-trial test, with Sidak’s multiple comparisons to compare trials 4 and 5. Each trial was also normalized relative to the first trial (baseline), and one-sample t-tests compared mean values of each trial to this baseline. For the three-chamber test, paired t-tests were used. For all these tests, null hypotheses were rejected at the 0.05 level.

## RESULTS

### C57Bl6/J mice can discriminate between siblings

The 5-trial test was used to assess the fine social discrimination abilities of 3- to 4-month-old female C57Bl6/J mice (Figure 1A). A significant reduction in interaction time with the familiar mouse was observed across trials, independent of the demonstrator’s sex (Trial effect: F(3,57)=17.79, p<0.0001; demonstrator sex effect: F(1,19)=1.004, p=0.3289, interaction: F(3,57)=0.191, p=0.9025; Two-way RM ANOVA, Figures 1B and 1C). When a sibling (brother or sister) of the mouse presented in trials 1-4 was introduced during trial 5, the C57Bl6/J females exhibited renewed interest, reflected by a significant increase in interaction time (Exploration time during trial 4 *vs* trial 5: female demonstrator, p<0.00019; male demonstrator, p<0.0001, Sidak’s multiple comparison test, Figure 1B), independent of the sex of the demonstrator. These results demonstrate that 3- to 4-month-old female C57Bl6/J mice are capable of finely discriminating between very similar mice, including siblings.

**Figure 1:**
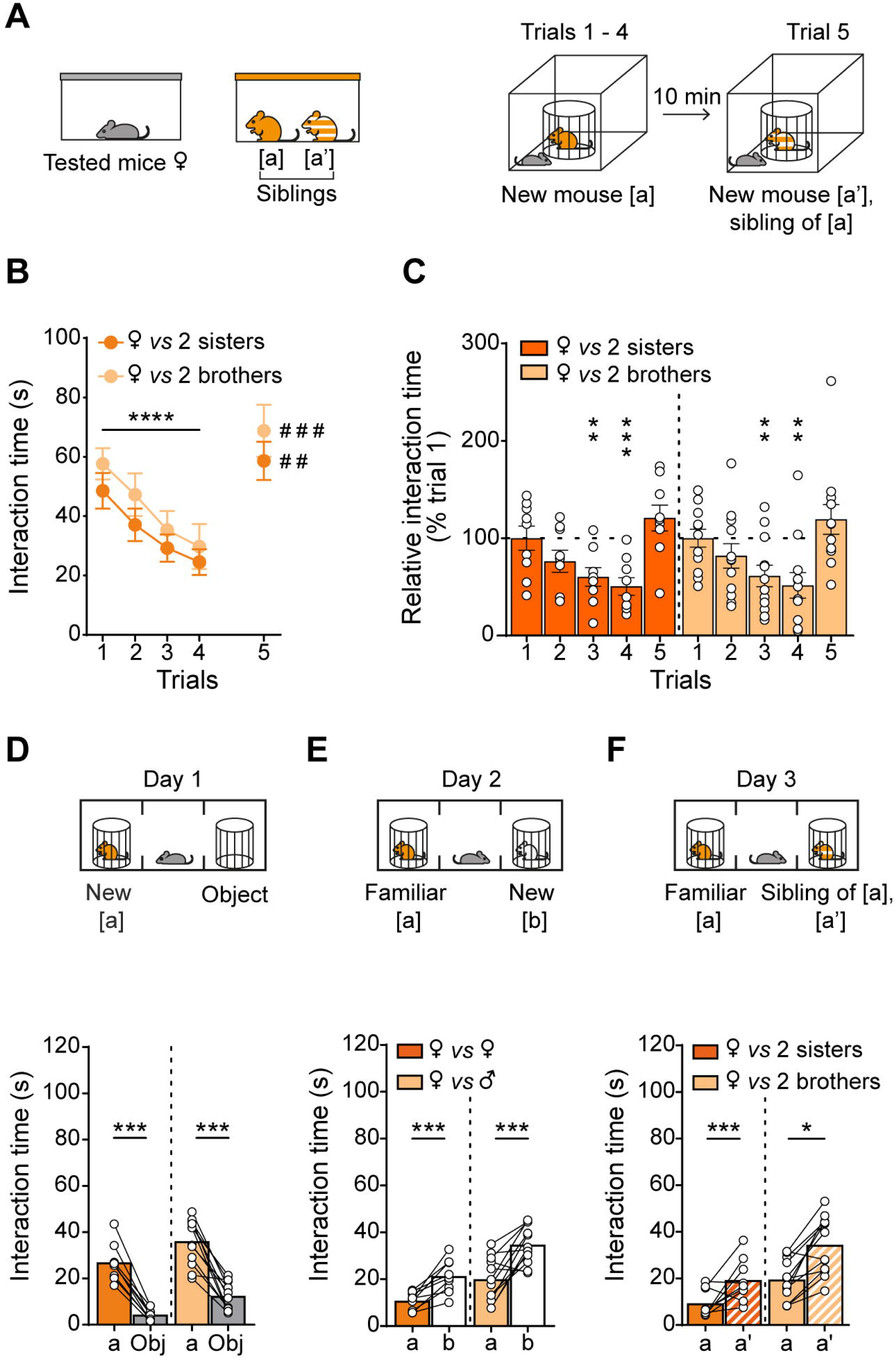
Adult female mice can discriminate between two siblings, regardless of their sex. (A) Paradigm to assess fine social discrimination. Tested mice are female, and demonstrator mice ([a] and [a’]) are either brothers or sisters from the same litter. During trials 1 to 4, the tested mouse is presented with a novel mouse [a]. On the last trial, [a’], a brother or sister of mouse [a], is presented. (B) Mouse interaction time in successive trials. Fine social memory assessed with the 5-trial test reveals that females can discriminate between two unrelated brothers (light orange) or sisters (dark orange) from the same litter. ***p<0.001, ^##^p<0.01, ^###^p<0.001, RM Two-way ANOVA followed by Sidak’s multiple comparisons. (C) Normalized measures show that interaction time decreases significantly during trials 1-4. One sample t-test **p<0.01 and ***p<0.001 between trial n and the baseline (100%). (D) Sociability test in the 3-chamber paradigm indicates that females spend more time with a conspecific, regardless of sex, than with an empty cage. ***p < 0.001 paired t-test between mouse and object condition. (E) The social discrimination test reveals that female mice spend more time with a new mouse than with the familiar one, ***p<0.01, paired t-test. (F) The fine social discrimination test reveals that female mice can discriminate between two siblings *p<0.005 and ***p<0.001 paired t-test. Bar graphs represent the mean and error bars SEM. Circles indicate individual animal values.

To test whether these fine social discrimination abilities persist over the long term, mice were subjected to a modified version of the three-chamber test (Figure 1D, E, F). On the final day (Figure 1F), female C57Bl6/J mice were tested for their ability to discriminate between two siblings (brothers or sisters). The tested mice showed a significant preference for the unfamiliar sibling (females *vs* females: n=9, familiar: 9.19±2,00 s, sibling: 19.11±3.03 s, paired-t-test: p=0.0197, t(8)=2.91, df=8; females *vs* males, n=12, familiar: 19.29±2.54 s, sibling : 34.22±3.59 s, paired t-test: p=0.0003, t(11)=5.16, df=11; Figure1F). These results suggest that fine discrimination between siblings relies on long-term social memory of individual features. A similar study conducted with male C57Bl6/J mice (Extended data, figureE1) confirmed that both male and female C57Bl6/J mice form robust short- and long-term social memories, allowing them to distinguish between closely related individuals regardless of their sex.

### Fine social discrimination in a mouse model of Alzheimer’s Disease (AD)

The Tg2576 mouse line is a well characterized model of AD that develops progressive, age-related memory deficits (Kosel et al., 2020). By 9 months of age, Tg2576 animals reach the symptomatic stage and exhibit impairments in social discrimination and social memory in the 3-chamber and 5-trial tests, respectively (Rey et al., 2022). In the present study, we investigated whether 3-month-old Tg2576 mice, *i.e.*, at pre-symptomatic stage, could perform fine social discriminate between closely related conspecifics. We postulated that fine social discrimination is more cognitively demanding than classic social discrimination, and hypothesized that it would be impaired at an earlier stage of pathology.

First, 3-month-old Tg2576 and NTg mice were subjected to the 5-trial test (Figure 2A). Both genotypes showed a significant reduction in interaction time from trials 1 to 4 (Trial effect: F(2.50,34.98)=32.41; p<0.0001; Genotype effect: F(1,14)=0.31; p=0.5860, Interaction: F(3,42)=1.83; p=0.1566; Two-way RM ANOVA, Figure 2B), suggesting that social learning is intact in youngTg2576 mice. When a novel, unrelated conspecific was introduced during trial 5, both NTg and Tg2576 mice exhibited renewed interest, reflected by an increase in interaction time (Exploration time during trial 4 vs trial 5: NTg, p=0.0302; Tg2576: p=0.0422, Sidak’s multiple comparison test, Figures 2B and 2C). These results indicate that 3-month-old Tg2576 mice retain intact social memory social discrimination abilities at this very early pathological stage.

**Figure 2:**
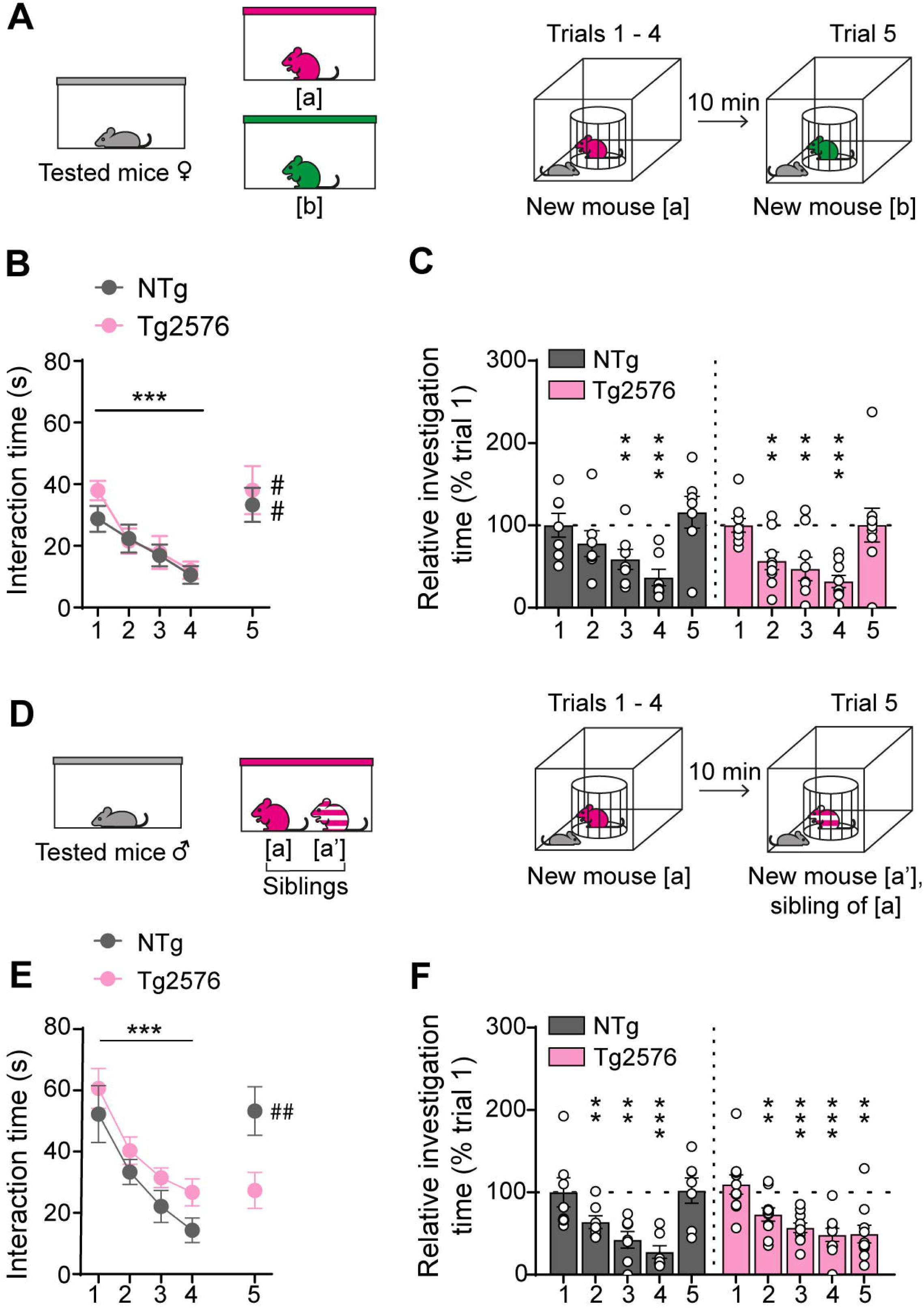
Deficits in fine social discrimination occur at preclinical stage in the Tg2576 mouse model of Alzheimer’s disease. (A) Paradigm to assess social discrimination. Tested mice are females Tg2576 and demonstrator mice ([a] and [b]) are unrelated female NTg mice from the same line. During trials 1 to 4, the tested mouse is presented with a novel mouse [a]. On the last trial, a novel mouse [b] is presented. (B) Interaction time during successive trials. The capacity to discriminate between two unrelated conspecifics is similar in Tg2576 (pink) and NTg (grey) mice, indicating intact social discrimination in Tg2576 mice at 3 months of age. ***p<0.05, ^#^p<0.05 RM Two-way ANOVA followed by Sidak’s multiple comparisons. (C) Normalized measures show that interaction time decreases significantly during trials 1-4 in both genotypes. One-sample t-test **p<0.01 and ***p<0.05 between trial n and the baseline (100%). (D) Fine social discrimination in Tg2576 mice is evaluated by presenting a novel mouse [a] (trials 1-4) followed by its sister [a’] (trial 5). (E) Mouse interaction time during successive trials. Fine social discrimination of two siblings is altered in Tg2576 (pink) compared to NTg (grey) mice, as indicated by the lack of interaction elicited by the presentation of the novel mouse [a’] (trial 5). ***p<0.05, ^##^p<0.01 Two-way ANOVA followed by Sidak’s multiple comparisons. (F) Normalized measures show that while interaction time decreases significantly during trials 1-4 in both genotypes, it also decreases when a novel but related mouse is presented to Tg2576 mice (pink). This indicates that fine social discrimination is altered as early as 3-month-old in Tg2576 mice. One sample t-test **p<0.01 and ***p<0.05 between trial n and the baseline (100%). Bar graphs represent the mean and error bars SEM. Circles indicate individual animal values.

Next, we assessed the ability of Tg2576 mice to distinguish between two closely related individuals (Figure 2D). To this end, mice were allowed to interact with two sisters in the same manner as described in Figures 1A and 1B. As expected, both NTg and Tg2576 mice recognized a novel individual across four successive trials (Trial effect: F(1.86,27.93)=34.64, p<0.0001; Genotype effect: F(1,15)=2.34, p=0.1470, Interaction: F(3,45)=0.181, p=0.9085; Two-way RM ANOVA, Figure 2E). However, when presented with a new sibling during trial 5, Tg2576 mice failed to show an increase in interaction time compared to trial 4, unlike NTg mice (Exploration time during trial 4 *vs* trial 5, NTg: p=0.0079, Tg2576: p>0.9999, Sidak’s multiple comparison test, Figures 2E and 2F). These findings suggest that while 3-month-old Tg2576 mice retain intact general social memory, they are unable to perform fine social discrimination, revealing an early cognitive impairment in this AD mouse model.

## DISCUSSION

This study provides new insights into the individual discrimination capacities of rodents. Indeed, our findings demonstrate that mice are capable of fine individual discrimination, a capacity that had, to our knowledge, not been previously reported. Furthermore, we hypothesized that fine social discrimination, a more nuanced form of social memory, would be affected earlier in the course of AD compared to broader social discrimination abilities. Given that the Tg2576 mouse model of AD exhibits altered social memory at 9 months of age (Rey et al., 2022), we tested fine social discrimination at younger, considered pre-symptomatic mice. Our findings reveal that although general social memory is intact, 3-month-old Tg2576 mice already exhibit impairments in fine social memory. This suggests that these two tasks, while related, are governed by different cognitive mechanisms.

The assessment of this novel form of social discrimination raises important questions: What role does fine discrimination play in natural settings? Does it hold ecological significance? What sensory cues enable such fine discrimination? What are the underlying neurological mechanisms, and why is this ability impaired early in Tg2576 mice? Could impaired fine social discrimination serve as an early indicator of AD?

### Role of fine social discrimination

Fine individual discrimination may be critical for maintaining social group stability and fitness, as it provides information on social status, relationships, fighting ability, territorial boundaries, and cooperation. By accurately identifying and recognizing the individuals, mice can adapt their behavior accordingly, thereby enhancing their chances of survival. This fine discrimination may also play a role in maintaining social hierarchy, allowing mice to recognize the dominance status of others and decide whether or not to engage in confrontations. This could reduce the likelihood of injury by minimizing unnecessary fights (Hare, 1987; Ligout and Porter, 2006). However, the neural networks underlying this fine social discrimination remains to be explored.

### Cues for fine social discrimination

Ligout & Porter (2006) have proposed several mechanisms for facilitate social discrimination, including indirect familiarization, where individuals learn cues from conspecifics as a reference for future encounters (Beecher, 1982). This includes both ”autoreference”, where individuals compare others to their own phenotypic traits (“self-matching”), and “alloreference”, where individuals use learned traits from familiar conspecifics for future recognition (“phenotype matching”). In the present case, we hypothesized that mice utilize phenotypic matching to perform fine social discrimination. This capacity to identify individuals based on comparisons with previously encountered conspecifics has also been observed in other rodents such as rats (Hepper, 1991) and squirrels (Holmes, 1986).

In addition, Beecher (1982) suggested the existence of individual signatures that enable social recognition. In mammals and in particular in rodents, recognition of these signatures likely involves multiple sensory modalities. While chemosensory cues (part of an olfactory signature) are often associated with social stimuli (Rodents: Camats Perna & Engelmann, 2015; Mammals: Sanchez-Andrade & Kendrick, 2009), other sensory modalities, such as vocalizations, also play a significant role in social communication and discrimination (Simola and Granon, 2019). In rodents, the frequency of vocalizations can vary based on the emotional context, reflecting the internal state of the sender and influencing the behavior of the listener (Brudzynski, 2013; Simola and Granon, 2019). Additionally, physical contact and tactile exploration, especially *via* whiskers during facial examination, are critical mediators of social interactions in mice (Arakawa, 2020). Indeed, mice with impaired whisker function due to whisker-trimming show decreased facial exploration during direct social interactions (Arakawa, 2020). Visual cues, such as tail rattling, freezing or grooming, can also convey information about a mouse’s emotional state (Jabarin et al., 2022). Notably, social discrimination is impaired when the social stimulus is anesthetized, suggesting that active behavior of social stimuli is required for successful social discrimination (de la Zerda et al., 2022). These multimodal sensory inputs are likely integrated to facilitate fine social discrimination, making it a complex process.

### Potential neuronal substrates of fine social discrimination

The CA2 area of the hippocampus plays a crucial role in social cognition in mice. Genetic inactivation of pyramidal cells in the dorsal part of CA2 has been shown to impair social memory (Hitti & Siegelbaum, 2014), suggesting that area CA2 might also contribute to fine social discrimination. Notably, anatomical changes implying modifications of neuronal excitability and plasticity are observed as early as 3 months of age in area CA2 of Tg2576 mice (Cattaud et al., 2018; Rey et al., 2022).

Recent studies have revealed that changes in adult hippocampal neurogenesis can also impact social memory. Adult-born granule neurons project to the CA2 area (Llorens-Martín et al., 2015), and this pathway is involved in distinguishing between novel and familiar individuals (Pereira-Caixeta et al., 2018; Cope et al., 2020). For example, reducing adult neurogenesis in the conditional TK transgenic mouse line leads to deficits in long-term social memory (Cope et al., 2020), while enhancing neurogenesis can rescue social memory deficits (Monteiro et al., 2014). Altogether, these findings indicate that the social aspects of episodic-like memories, such as the ability to form integrated memory of “what, where, and when” a conspecific was encountered, are particularly sensitive to changes in adult hippocampal neurogenesis. We have previously reported that adult hippocampal neurogenesis is impaired in 3 month-old Tg2576 mice (Krezymon et al., 2013). This, coupled with the fine social memory deficits observed in the present study, prompt us to postulate that these cognitive impairments may be linked to disrupted adult neurogenesis.

Furthermore, we propose that fine social discrimination, in particular the ability to distinguish between closely related individuals, may involve a process akin to pattern separation, where similar inputs are transformed into distinct neuronal representations (Yassa and Stark, 2011). Based on computational models and experimental data, adult-born hippocampal neurons are thought to play a key role in this process (Aimone et al., 2011; Sahay et al., 2011b). The elimination or reduction of adult-born neurons impairs pattern separation capacity (Clelland et al., 2009; Tronel et al., 2012), while increasing adult neurogenesis enhances the ability to distinguish between highly similar inputs (Sahay et al., 2011b, 2011a). Thus, we hypothesize that fine social discrimination may represent a form of “social pattern separation”. The behavioral paradigm we have developed in the present study could serve as a valuable tool for assessing social pattern separation abilities in mice.

### Fine discrimination deficits as a potential marker of early AD

Our findings show that impairments in fine social discrimination occurs early in AD pathology, in a mouse model of the disease, even before other social memory deficits become apparent. These deficits could serve as an early marker of the disease. AD patients experience a deterioration of daily activities due to cognitive and functional decline (Tarawneh and Holtzman, 2012). Identifying early signs of the disease and understanding the underlying mechanisms are major challenges. According to Rankin et al. (2008), changes in social behavior are often among the first signs of neurodegenerative diseases. Additionally, our data are consistent with findings from Pietropaolo et al. (2012), who reported that Tg2576 females exhibit an age-related decline in social investigation.

Understanding the mechanisms underlying fine social memory could inform strategies to prevent or delay social cognitive deficits. For humans, social activity and enriched social networks can help prevent these deficits (Qiu et al., 2009). Similarly, modulating the social environment of mice might delay the onset of fine social deficits in AD models. Research on rats has shown the importance of a social enrichment for social memory performance (Toyoshima et al., 2018) and environmental enrichment by modulating social plasticity, enables rodents to optimize their social relationships (Gubert and Hannan, 2019).

### Conclusion

This study highlights a previously uncharacterized form of social discrimination in mice—fine social discrimination—and demonstrates its impairment at early stage of AD in a mouse model. These findings strongly suggest that fine social deficits could serve as early markers of AD pathology. However, several key questions remain, such as the specific cues and neural substrates involved in this process, thus calling for further research.

## Conflict of interest

The authors declare no competing financial interests

## Acknowledgments

The authors thank Maud Combe for developing the Ethoc software. The mice were housed in the ABC Facility of ANEXPLO, Toulouse. The authors gratefully acknowledge the Mouse Behavioral Core (MBC) of the Center of Integrative Biology. This work was supported by the Centre National de la Recherche Scientifique (CNRS), the University of Toulouse, the Association France Alzheimer (AAP SM 2018 #1823), the Fondation Vaincre Alzheimer (Grant #61144). L.M.P.F. received a PhD fellowship from the French Ministry of Research.

---

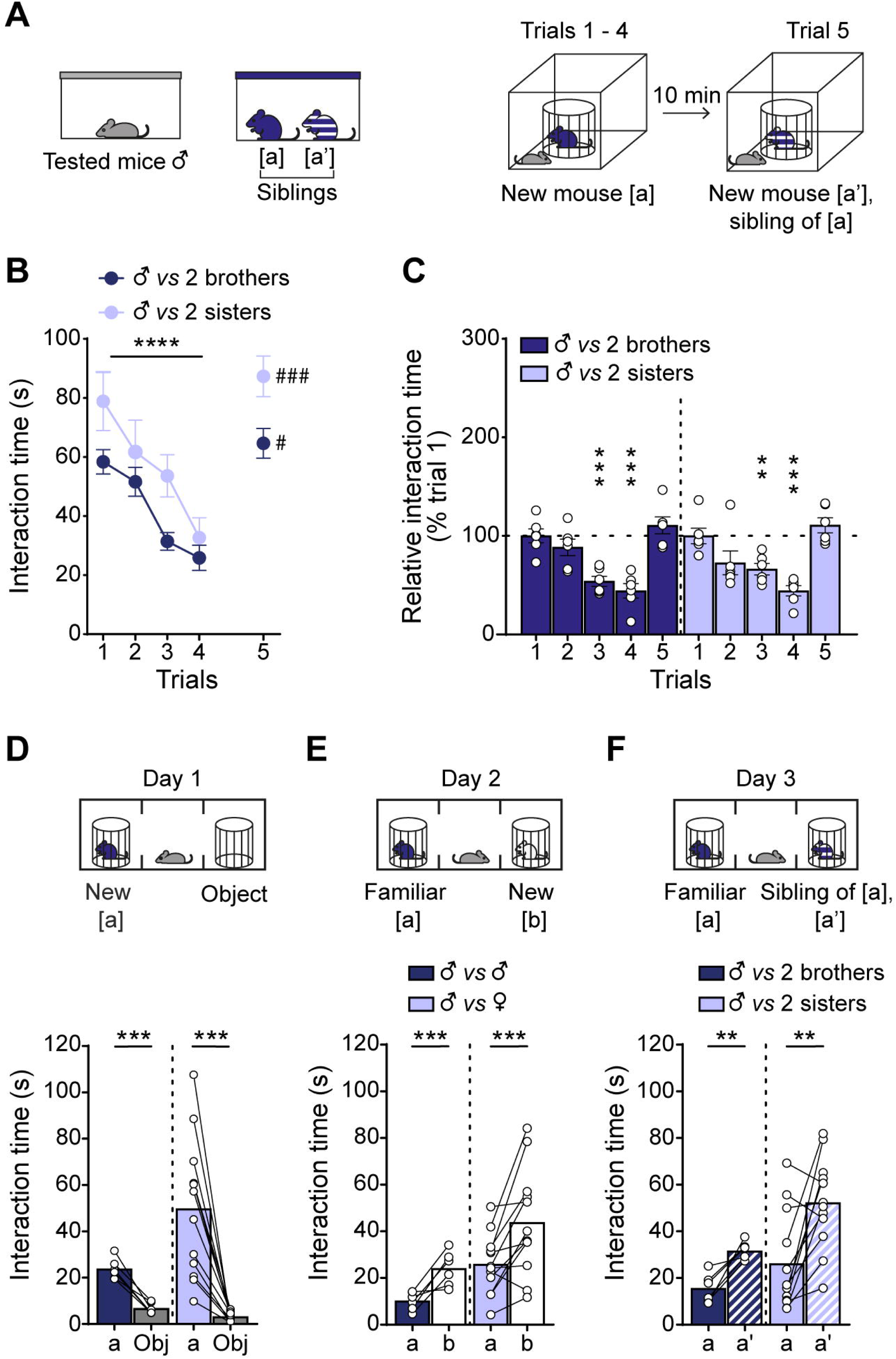

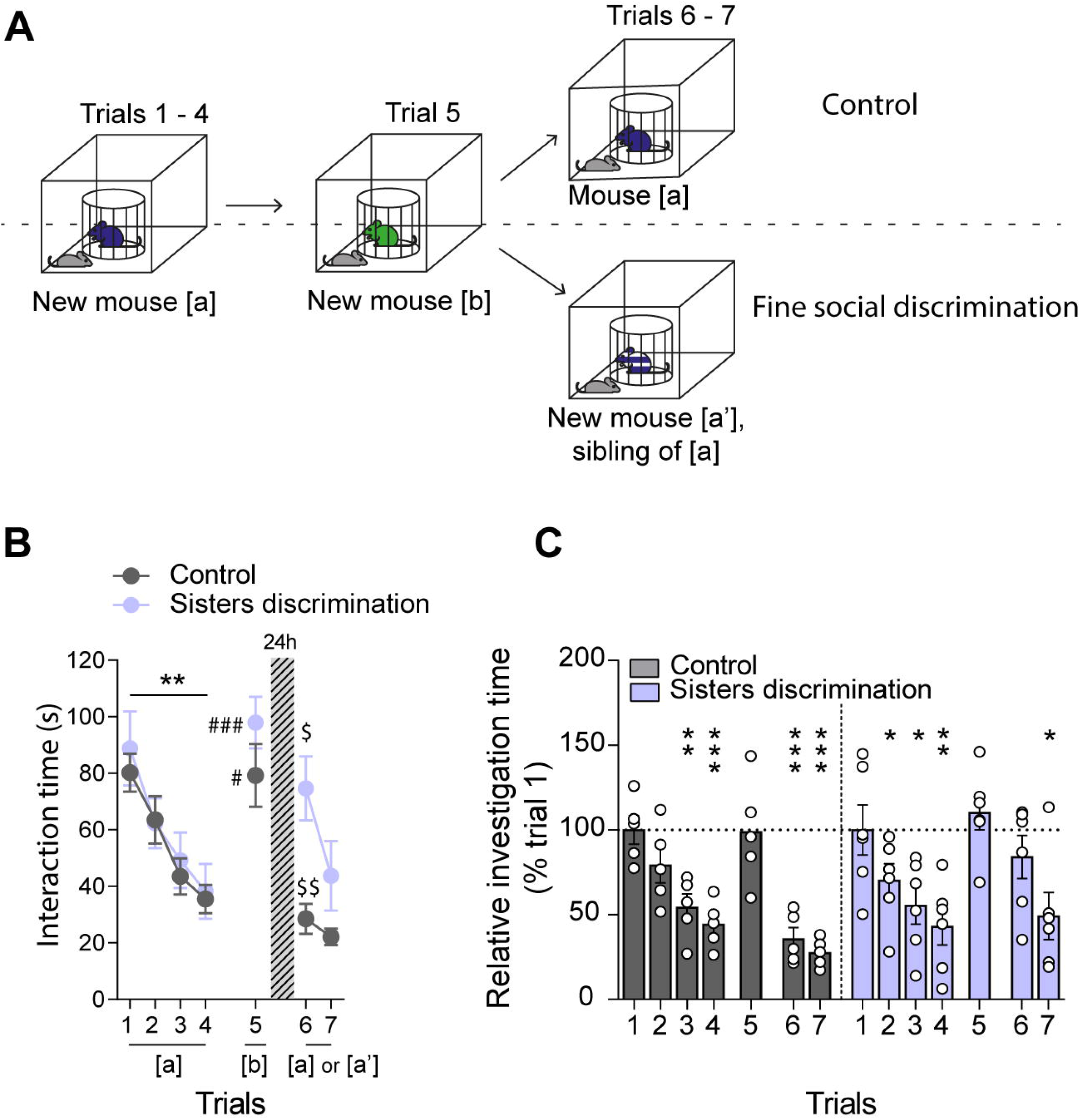

## Notes

### Competing Interest Statement

The authors have declared no competing interest.

