## Extended Figure Legends for "Fine Social Discrimination of Siblings in Mice: Implications for Early Detection of Alzheimer’s Disease"

EXTENDED DATA

**Fig E1: Adult male mice can discriminate between two siblings regardless of their sex**

(A) Paradigm for evaluating fine social discrimination. Tested mice are male, and demonstrator mice ([a] and [a']) are either brothers or sisters from the same litter. During trials 1 to 4, the tested mouse is presented with a novel mouse [a]. On the last trial, [a'], a brother or sister of mouse [a], is presented.
(B) Mouse interaction time in successive trials. Fine social memory, assessed with the 5-trials test, reveals that male mice can discriminate between two unrelated brothers (dark blue) or sisters (light blue) from the same litter.  ***p<0.05, ^#^p<0.05, ^###^p<0.001, RM Two-way ANOVA followed by Sidak’s multiple comparisons.
(C) Normalized measures show that interaction time decreases significantly during trials 1-4. One sample t-test **p<0.01 and ***p<0.001 between trial n and the baseline (100%).
(D) Sociability test in the 3-chamber paradigm indicates that male mice spend more time with a conspecific, regardless of sex, than with an empty cage. ***p<0.001, paired t-test between mouse and object conditions.
(E) The social recognition test reveals that male mice spend more time with a new mouse than with the familiar one, ***p<0.01, paired t-test.
(F) The fine social recognition test reveals that male mice can discriminate between two related individuals **p<0.01 and ***p<0.001, paired t-test.

Bar graphs represent the mean, and error bars represent SEM. Circles indicate individual animal values.

**Fig E2:** **Male mice are able to form a long-term social memory of two siblings.**

(A) Paradigm for evaluating long-term fine social memory of siblings. Tested mice are male, and demonstrator mice are female. During trials 1 to 4, the tested mouse is presented with a novel mouse [a]. In the following trial (trial 5), a novel mouse [b], unrelated to [a], is presented. After 24 hours, the same mouse [a] (control group) or its sister [a’] (fine social discrimination) is presented during the following two trials (6-7).
(B) Mouse interaction time over successive trials. In both groups, investigation time decreases significantly from trial 1 to 4, **p<0.01, RM Two-way ANOVA. The presentation of a novel individual triggers a rebound in interaction in both groups (trial 5), ^#^p<0.05, ^###^p<0.001, paired t-test T4 *vs* T5. On the next day (trials 6 -7), when a familiar individual is presented, a reduced interaction time is observed compared to trial 1 and trial 5, ^$$^p<0.01, paired t-test T5 *vs* T6. However, when the sister of the familiar individual is presented, the interaction time remains high and similar to trial 1.
(C) Normalized interaction time increases significantly at trial 6 in presence of the sister of the mouse presented the day before (Trial 1 *vs* Trial n). One sample t-test **p<0.01 and ***p<0.001 between trial n and the baseline (100%).

Bar graphs represent the mean, and error bars represent SEM. Circles indicate individual animal values.
